# Vesicle Internalization Proceeds via a Morphological Phase Transition

**DOI:** 10.64898/2026.06.16.732530

**Authors:** Itay Schachter, Pavel Jungwirth, Daniel Harries

## Abstract

Vesicle internalization proceeds through a series of multivesicular topologies essential for endocytic transport and cellular compartmentalization. The energetic landscapes of related transitions, including vesicle budding and pearling, are known to be governed by the coupling of spontaneous curvature, leaflet area asymmetry, and reduced volume. However, the physical principles driving the structural transformation of hemifused intermediates remain unresolved. Using a continuum elastic model, we identify a morphological phase transition in hemifused invaginating vesicles, from an initial lens-like geometry to an elongated “kettle” geometry. This transition is discontinuous as long as the invaginating vesicle’s reduced volume is below a critical threshold, but continuous otherwise. The kettle-like morphology is metastable across a broad range of leaflet area asymmetries, potentially enabling a hysteretic externalization pathway. Increasing either the spontaneous curvature of the shared outer leaflet or the size of the invaginating vesicle, alone or in tandem with the host vesicle, turns the kettle morphology into the global free energy minimum. Notably, simply scaling up the size of both vesicles does not eliminate the free energy barrier. This quantitative characterization provides a structural reference for identifying internalization intermediates witnessed in experimental imaging, and maps the morphological evolution of the internalization pathway across its physical parameter space.

## 1 Introduction

Vesicle internalization is a necessary mechanical precursor to endocytic transport and cellular compartmentalization. ^1,2^ This internalization process involves the physical translocation of the invaginating vesicle volume across the hemifusion junction plane into the host lumen. The energetic landscapes of simple morphological transitions—such as budding, pearling, or fission—have been extensively analyzed in terms of lipid spontaneous curvature and leaflet area asymmetry ^3–8^. However, the physical principles governing the internalization of hemifused intermediates and their associated energetic landscape remain unresolved. Specifically, the internalization process is likely not merely a geometric reconfiguration; instead, it involves a complex partitioning of elastic strain across shared membrane leaflets^3^. Understanding the forces that drive vesicle internalization from the exterior to the interior of a host vesicle is critical for deciphering how cells regulate cargo uptake at the nanoscale.

Traditional theoretical frameworks, such as the area-difference elasticity model, ^3^ have successfully described shape transitions in isolated vesicles by assuming that membrane thickness is negligible compared with the vesicle radius. However, these models struggle with the topological complexity of hemifused vesicles, where the shared outer leaflet between the two vesicles and distinct inner leaflets impose coupled constraints on leaflet area and volume. Furthermore, at the nanoscale, the non-negligible membrane thickness and the high curvatures required for neck constriction introduce energetic penalties that simple continuum models often neglect ^9,10^. To date, the pathway through which an invaginating vesicle evolves into a fully internalized vesicle has lacked a quantitative energetic description.

Recent cryo-electron tomography observations have identified specialized vesicular complexes, termed ‘hemifusomes’, which exhibit a continuum of intermediate morphologies ranging from shallow lens-like geometries to deep, elongated invaginations. ^11^ While these structures suggest an active, regulated internalization pathway—potentially steered by an associated proteolipid nanodroplet— the underlying mechanical and energetic landscapes of these transitions remain largely conjectural.

In this work, we use a detailed continuum elastic model, resolved using a finite-element solver, to map the internalization pathway of hemifused large unilamellar vesicles (LUVs). We identify a previously uncharacterized morphological phase transition from a lens-like geometry to an elongated “kettle” geometry. We show that this transition is regulated by the reduced volume of the invaginating vesicle, and manifests as either a discontinuous, hysteretic snap-through transition or by a continuous structural evolution. By varying physical parameters, including lipid spontaneous curvature and system scale (preserving relative vesicle dimensions), we demonstrate that this transition acts as a binary mechanical switch between morphologies. Our findings provide a structural reference for interpreting experimental imaging of associating vesicles and offer a physical mechanism by which specialized biological lipidic complexes bypass kinetic traps to achieve efficient internalization.

## 2 Theoretical Model

Hemifused vesicles are modeled via elastic surfaces based on the Helfrich–Hamm–Kozlov formalism ^9,10,12^. We utilize the numerical implementation of a Finite-Element solver^6,13^ detailed in our previous work ^14^. The current version incorporates subtle differences to the grid discretization, parameter space, and minimal free energy path sampling, as described in the SI. The model treats the two inner- and shared outer-leaflets as distinct slabs connected via midplane surfaces, Fig. 1. The total free energy *F* for each leaflet *j* is given by:

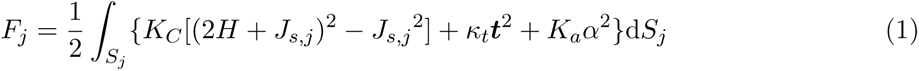

**Figure 1:**
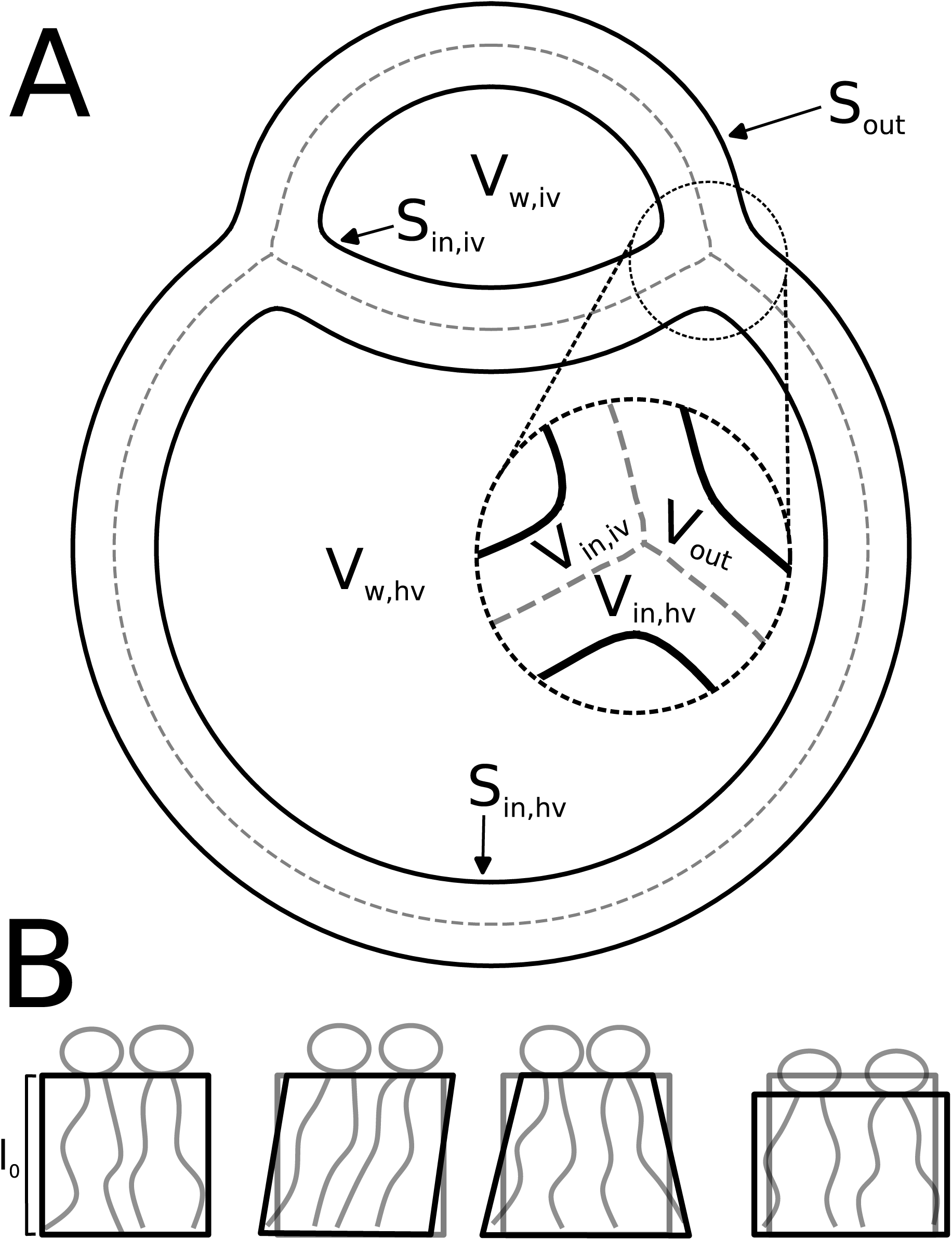
(A) Cross-sectional schematic of two hemifused vesicles defining leaflet surfaces (*S*_out_*, S*_in,iv_*, S*_in,hv_), midplane surfaces (dashed lines), and entrapped water volumes (*V*_w,iv_*, V*_w,hv_). The labels within the bilayer denote the shared outer (*V*_out_) and individual inner (*V*_in,iv_*, V*_in,hv_) leaflet volumes. (B) Deformation modes considered in the elastic model (left to right): undeformed state with relaxed lipid length *l*_0_, tilt, bend, and _4_area-compressibility.

The first term in the integral accounts for the bending energy. The bending modulus *K_C_* accounts for the energetic penalty for splay deformations, 2*H* = ∇ · ***n***, relative to (minus) the spontaneous curvature *J_s,j_*of leaflet *j* (with curvature defined as positive for spherical micelles). The second term describes the tilt energy, regulated by the tilt modulus *κ_t_* and the tilt vector ***t*** = ***n****/*(***N*** · ***n***) − ***N*** , with ***N*** representing the surface normal. The third term characterizes the area-compressibility energy, with modulus *K_a_* accounting for fluctuations in the local area-per-lipid, *a*, from its equilibrium value *a*_0_, via the relation 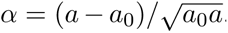. Under the assumption of local lipid incompressibility, *al* = *a*_0_*l*_0_, this expression simplifies to 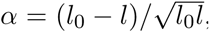, where *l* (*l*_0_) is the lipid (relaxed lipid) length.

To maintain consistency with typical experimental conditions, the model enforces the following volumetric constraints. Each leaflet’s volume is held constant, reflecting global lipid incompressibility and the negligible rate of phospholipid flip-flop ^15^. Similarly, the volumes of the entrapped water, *V*_iv_ and *V*_hv_, are fixed; these remain effectively invariant under specific osmotic conditions ^4^ and can be determined by the vesicle fabrication method^16^. The implementation code is hosted on GitHub.

Vesicle internalization geometrically requires changing the leaflets volumes. Consider two initially spherical vesicles, termed the host (HV) and invaginating (IV) vesicles, each with midplane radii of *R*_m,hv_ = 50 nm and *R*_m,iv_ = 15 nm. If the two vesicles are connected by an infinitely thin fusion stalk but the IV is not yet invaginated, the configuration is referred to as complete outer stalk. The ideal volume of the shared outer leaflet is 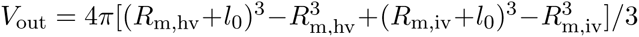 and the HV inner leaflet’s volume is 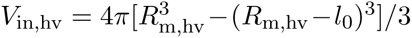. We fix the IV inner leaflet’s volume to 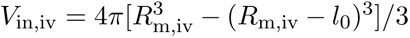 which is its ideal volume as a vesicle of radius *R*_m,hv_. We can further define a complete inner stalk state, where the invaginating vesicle has invaginated and is only connected to the host vesicle by an infinitely thin fusion stalk. In this state (neglecting entrapped water for the internalization coordinate derivation) 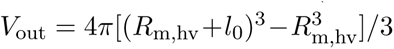 and 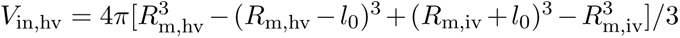. To follow the internalization process, we define an internalization coordination *p*_0_ such that *V*_1_ and *V*_2_ are interpolated linearly between the complete outer (*p*_0_ = 1) and inner (*p*_0_ = −1) stalk states. Finally, the entrapped water volumes, *V*_w,hv_ and *V*_w,iv_, may deviate from their ideal values of 4*π*(*R*_m,hv_ − *l*_0_)^3^*/*3 and 4*π*(*R*_m,iv_ − *l*_0_)^3^*/*3 respectively, with *ν*_hv_ and *ν*_iv_ defined as the relative water content factors; by default, *ν*_iv_ is set to 1 unless specified otherwise.

The default lipid model parameters for the subsequent analyses correspond to determined values for 1-palmitoyl-2-oleoyl-*sn*-glycero-3-phosphocholine (POPC) at *T* = 298.15 K and are defined as follows ^17,18^: *K_C_* = 14.49 *k*_B_*T* , *κ_t_* = 9.5 *k* _B_*T* · nm^−2^, *K_a_* = 28.93 *k* _B_*T* · nm^−2^, and *l*_0_ = 1.51 nm. Since the spontaneous curvature of POPC is negligible ^19^, *J_s_* is set to zero. Introducing these parameters into the model enables the investigation of internalization under experimentally relevant conditions. We focus on structures where the host vesicle has a prolate morphology, which follows the experimentally observed shape in hemifusomes ^11^ and is further discussed in the next section.

## 3 Results

### 3.1 Internalization Bifurcates into Discrete “Eye” and “Kettle” Morphological Phases

We mapped the morphological landscape of the hemifused complex by minimizing the free energy *F* (Eq. 1) with respect to the internalization coordinate *p*_0_ ∈ [−0.8, 0.4], while maintaining a fixed invaginating vesicle (IV) reduced volume of *ν*_iv_ = 0.75. This minimization identifies two distinct equilibrium branches: the “eye” and “kettle” morphologies (Fig. 2).

**Figure 2:**
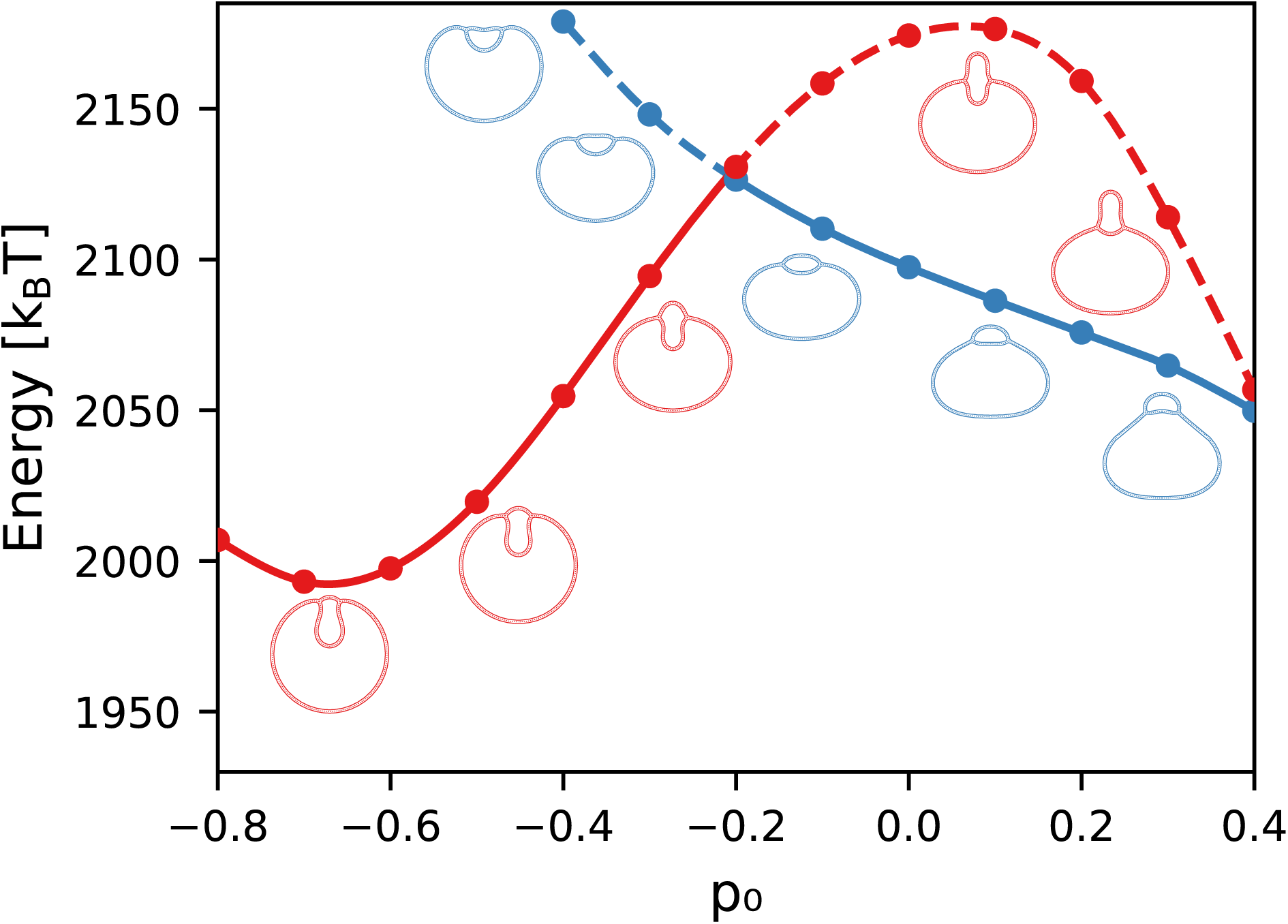
Energy landscape of the “eye” (blue) and “kettle” (red) morphologies as a function of the internalization coordinate *p*_0_ at *ν*_iv_ = 0.75. Scatter points indicate discrete *p*_0_ coordinates of the energy minimizations, lines are cubic spline interpolation of these points. Full lines indicate the global minimum free energy, equilibrium state, which exhibits a sharp cusp at the transition point. Dashed lines represent metastable branches. This cusp marks the discontinuous energy derivative which is a signature of a first-order jump. Representative structures for each morphology are indicated along the internalization path.

The “eye” morphology, characterized by a lens-shaped IV that maintains a wide hemifusion neck, constitutes the equilibrium structure for *p*_0_ *>* −0.211. Conversely, the “kettle” morphology—defined by an elongated IV and a constricted neck—becomes the stable phase at lower *p*_0_ values. As *p*_0_ decreases toward the complete inner stalk state (*p*_0_ = −1), both morphologies exhibit a progressive increase in the internalization ratio (Φ_in_), defined as the fraction of the IV volume residing on the HV side of the hemifusion junction plane.

A notable feature of this energy landscape is the broad range of metastability, where both morphologies persist as local minima beyond their respective stability limits. This is particularly pronounced for the kettle morphology, which remains metastable across the entire investigated range of *p*_0_. While the kettle state again becomes thermodynamically favored over the eye state at *p*_0_ ≈ 0.41, this regime concomitantly involves a transition of the HV into an oblate “eye” geometry for *p*_0_ ≳ 0.1 (see Fig. S1). Since such oblate deformations have not been observed in experimental studies of biologically hemifused vesicles ^11^, we focus here on the physiologically relevant prolate and spherical HV configurations.

### 3.2 The First-Order “Eye”–“Kettle” Transition Terminates at a Critical *ν*_iv_

The transition from the “eye” to “kettle” morphology is regulated by the reduced volume of the invaginating vesicle, *ν*_iv_. At low *ν*_iv_, the energy landscape exhibits a sharp cusp at the intersection of the two branches, signaling a discontinuous derivative of the free energy and a morphological first-order transition jump (Fig. 3). As *ν*_iv_ increases, this metastability region narrows, indicating the approach toward a critical point located between *ν*_iv_ ≈ 0.825 and 0.85. Beyond this critical threshold, the cusp vanishes, and the system follows a continuous structural continuum. This smooth evolution is visually confirmed at *ν*_iv_ = 0.875 (Fig. 4), where the IV lens elongates without undergoing a discrete jump in neck constriction or aspect ratio.

**Figure 3:**
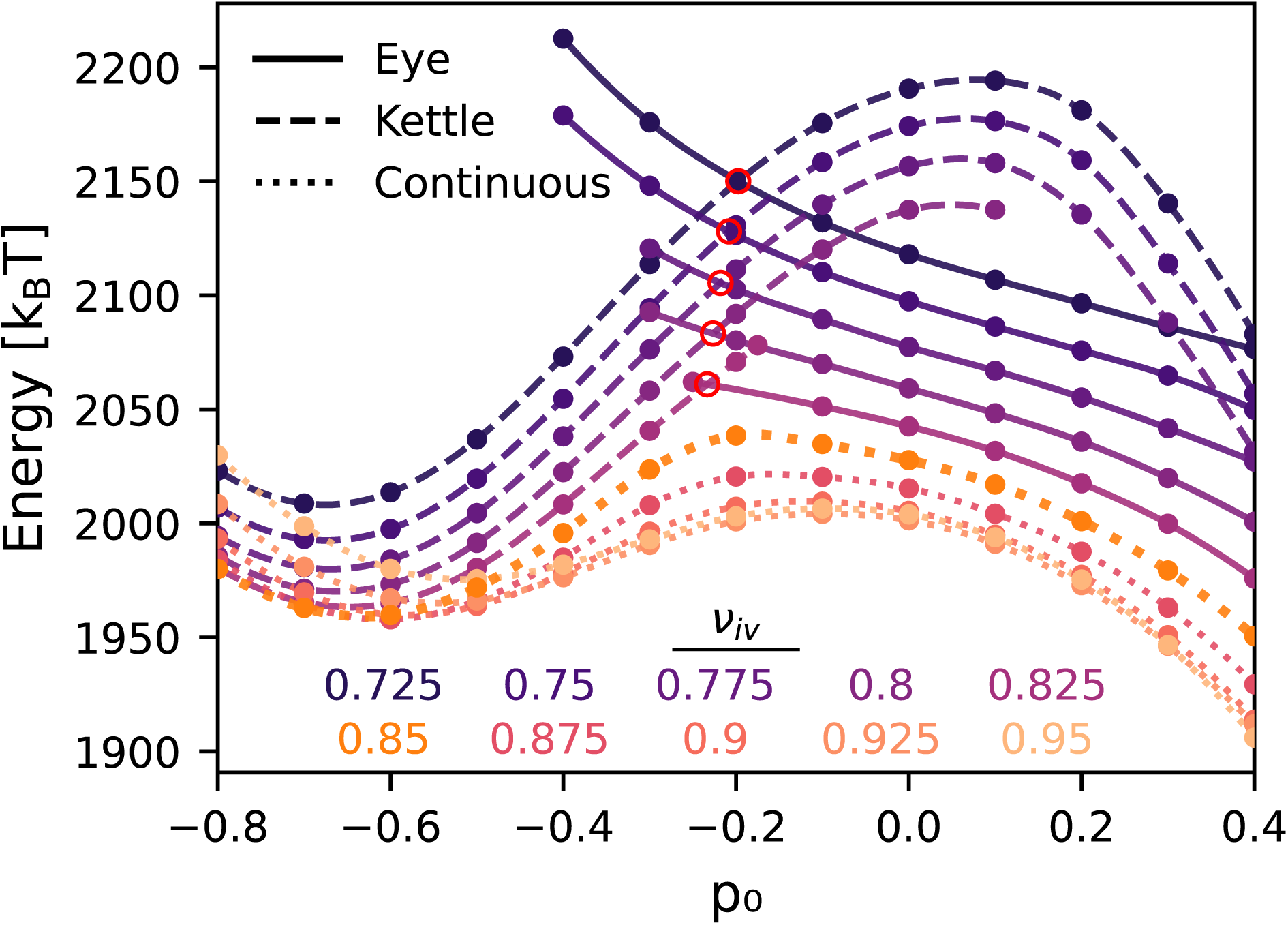
Energy landscape as a function of the internalization coordinate *p*_0_ for varying reduced volumes *ν*_iv_ ∈ [0.725, 0.95]. Scatter points indicate discrete *p*_0_ coordinates from the energy minimization protocol; lines are cubic spline interpolations. Solid lines represent the eye morphology, dashed lines represent the kettle morphology, and dotted lines indicate the continuous regime. Red open circles mark the thermodynamic crossover points where the eye and kettle branches are isoenergetic (*E*_eye_ = *E*_kettle_). As *ν*_iv_ increases (transitioning from dark to light curves), the energy barrier between morphologies decreases and the metastability regions shrink. Beyond a critical volume (*ν*_iv_ ≳ 0.85), the discontinuous bifurcation vanishes, and the system follows a single continuous path toward internalization.

**Figure 4:**
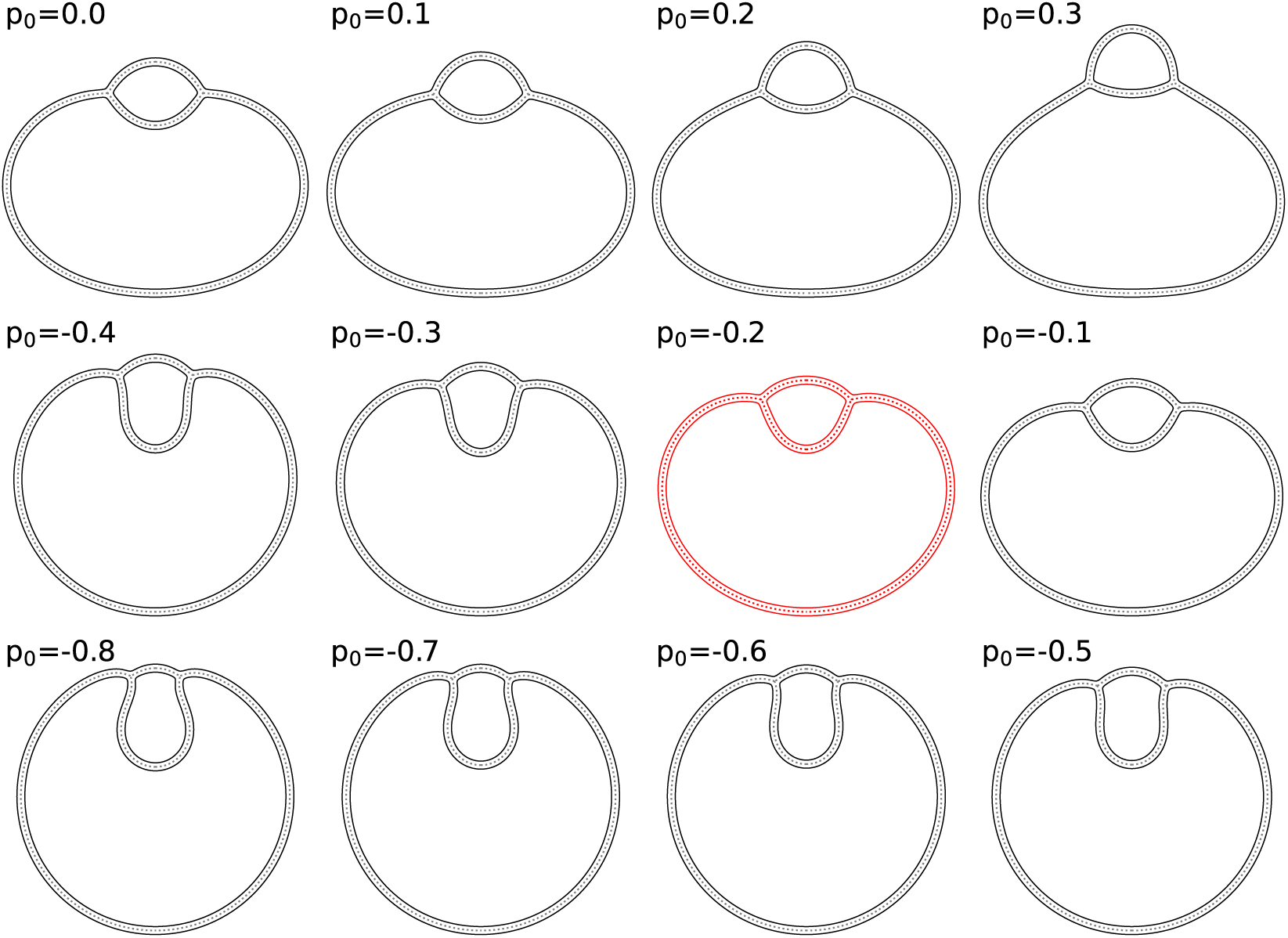
Axisymmetric cross-sections illustrating the continuous internalization pathway for *ν*_iv_ = 0.875. The sequence follows the structural evolution across *p*_0_ [ 0.8, 0.3], depicting the smooth deformation from an external protrusion (top row) to a deep invagination (bottom row). The profile at *p*_0_ = − 0.2 (highlighted in red) corresponds to the intermediate configuration near the crossover point between the eye and kettle morphologies seen in more deflated invaginating vesicles.Figure 5: Phase diagrams for internalized vesicle reduced volumes *ν*_iv_ of 0.725, 0.75, 0.775, 0.8, 0.825, and 0.85 (panels A–F, respectively) as a function of host vesicle reduced volume *ν*_hv_ and internalization coordinate *p*_0_. The solid black line indicates the thermodynamic transition between eye and kettle morphologies, which occurs for *ν*_iv_ *<* 0.85. Both the transition boundary and the continuous color map representing the internalization ratio Φ_in_ were generated via cubic spline interpolation of the discrete simulation data. The vanishing of the Φ_in_ discontinuity across the boundary illustrates the structural convergence as the system approaches the critical point in panel F.

We mapped the thermodynamic stability of both morphologies across host vesicle reduced volumes *ν*_hv_ ∈ {0.85, 0.9, 0.95, 1.0} and internalization coordinate *p*_0_ ∈ {−0.5, −0.4*, . . . ,* 0.4} (Fig. 5). The transition boundary, representing the coexistence locus where the eye and kettle states swap global stability, remains robust across the entire investigated range of *ν*_hv_. As the HV swells and approaches a spherical geometry (*ν*_hv_ → 1.0), the transition line shifts toward higher *p*_0_; conversely, HV deflation slightly broadens the stability range of the eye morphology.

**Figure 5:**
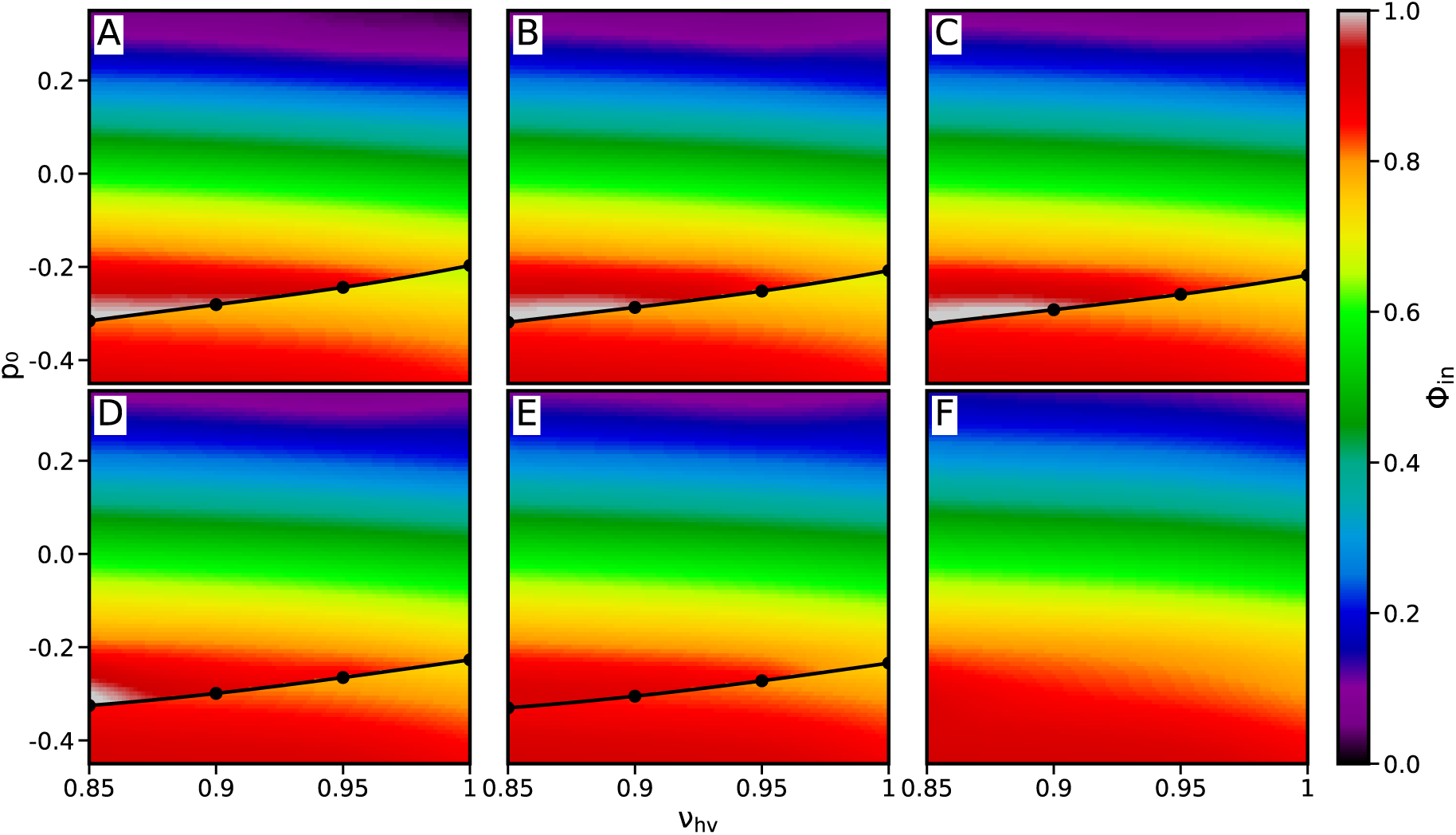
Phase diagrams for internalized vesicle reduced volumes *ν*_iv_ of 0.725, 0.75, 0.775, 0.8, 0.825, and 0.85 (panels A–F, respectively) as a function of host vesicle reduced volume *ν*_hv_ and internalization coordinate *p*_0_. The solid black line indicates the thermodynamic transition between eye and kettle morphologies, which occurs for *ν*_iv_ *<* 0.85. Both the transition boundary and the continuous color map representing the internalization ratio Φ_in_ were generated via cubic spline interpolation of the discrete simulation data. The vanishing of the Φ_in_ discontinuity across the boundary illustrates the structural convergence as the system approaches the critical point in panel F.

The structural divergence between phases is quantified by the internalization ratio Φ_in_. In the discontinuous regime (*ν*_iv_ ≤ 0.825), Φ_in_ jumps sharply across the transition boundary. Internalization slightly retracts upon transitioning into the kettle morphology; Φ_in_ decreases abruptly at the crossover point before resuming its increase as *p*_0_ drops. However, the magnitude of this jump diminishes as *ν*_iv_ approaches the critical point, reflecting the vanishing structural distinction between the two morphological phases.

### 3.3 Energy Barriers for “Eye” and “Kettle” Transitions Scale Inversely with *ν*_iv_

To identify how the IV reduced volume (*ν*_iv_) regulates transition kinetics, we calculated the free energy barriers along the minimal free energy path (Fig. 6). Across all investigated configurations, barrier heights for both eye-to-kettle and kettle-to-eye transitions scale inversely with *ν*_iv_. In highly deflated vesicles (*ν*_iv_ = 0.725, Fig. 6, A), the kettle-to-eye barrier exceeds 175 k_B_T at *p*_0_ = −0.4 and remains above 70 k_B_T at the thermodynamic crossover. The energy barrier remains above 30 k_B_T throughout the inspected *p*_0_ range.

**Figure 6:**
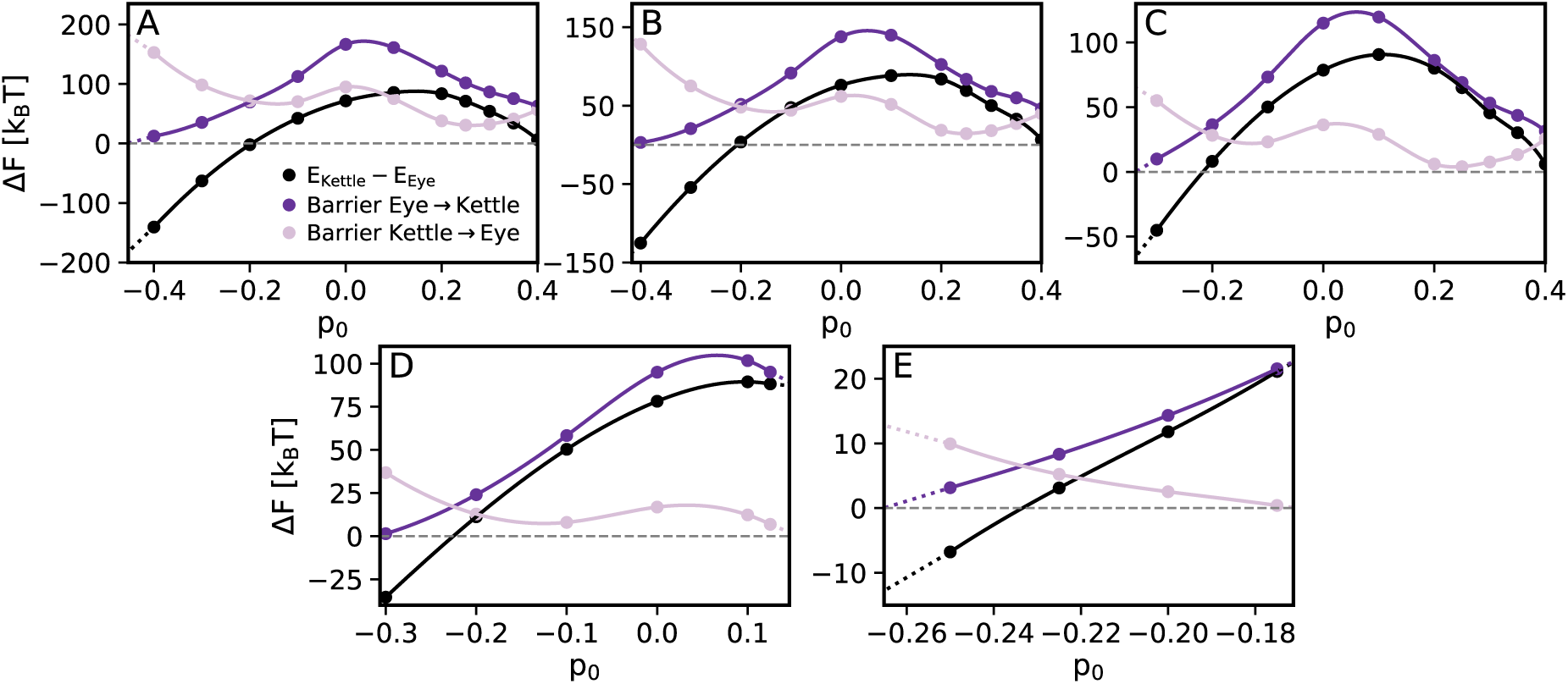
Free energy differences and kinetic barriers along the minimal free energy path between the “eye” and “kettle” morphologies for (A) *ν*_iv_ = 0.725, (B) *ν*_iv_ = 0.75, (C) *ν*_iv_ = 0.775, (D) *ν*_iv_ = 0.8 and (E) *ν*_iv_ = 0.825. The black curve denotes the energy difference between the “kettle” and “eye” morphologies Δ*F* = *E*_kettle_ *E*_eye_. We observe the thermodynamic crossover when Δ*F* crosses 0. Purple and light-purple curves represent the energy barriers for the eye-to-kettle and kettle-to-eye transitions, respectively. Scatter points indicate discrete *p*_0_ coordinates of the energy minimizations; solid lines represent cubic spline interpolations. The vanishing of these barriers marks the loss of metastability and signals the system’s approach toward the critical point in *ν*_iv_ ≈ 0.85.

Increasing *ν*_iv_ systematically destabilizes the metastable morphologies by shrinking the coordinate range where local minima persist. At low *ν*_iv_, the kettle-to-eye barrier fluctuates non-monotonically and maintains two local minima. As the vesicle swells, the landscape flattens and these minima vanish, which effectively truncates the kettle phase metastability. Specifically, at *ν*_iv_ = 0.775, the barrier vanishes near *p*_0_ = 0.2. As *ν*_iv_ reaches 0.8, this instability point shifts to *p*_0_ ≈ 0.15. By *ν*_iv_ = 0.825, the kettle morphology loses its local stability entirely around *p*_0_ = −0.15.

Increasing *ν*_iv_ abruptly truncates the kettle morphology’s metastability region at two discrete internalization coordinate thresholds. At *ν*_iv_ = 0.725, the kettle-to-eye barrier (pink line, Fig. 6) displays two distinct local minima at *p*_0_ ≈ 0.25 and *p*_0_ ≈ −0.15. These minima vanish sequentially as the vesicle swells, systematically cutting the kettle phase’s stability range. The first minimum nearly reaches zero at *ν*_iv_ = 0.775 and disappears entirely by *ν*_iv_ = 0.8, rendering the kettle state unstable at high *p*_0_. Increasing *ν*_iv_ further triggers another truncation of the metastability region at the second internalization coordinate; by *ν*_iv_ = 0.825, the kettle morphology loses its local stability at *p*_0_ ≳ −0.15, effectively shrinking the coexistence region from both ends of the internalization path.

The energy landscapes exhibit a distinct asymmetry in the morphological phase metastability regimes: the eye morphology stability at low *p*_0_ is significantly more limited than that of the kettle morphology at higher *p*_0_. As *ν*_iv_ approaches the critical point, the *p*_0_ range where both morphologies coexist narrows. This suggests that while deflated IVs can remain kinetically trapped in metastable states across a broad range of conditions, more swollen IVs are forced to transition within a significantly tighter coordinate window.

### 3.4 The “Kettle” Morphology Stability Depends on Vesicle Physical Properties

The stability of the kettle morphology was further evaluated against variations in IV radius (*R*_m,iv_), global system scale (*S*), and the spontaneous curvatures of the IV inner leaflet (*J_s,_*_iv_) and shared outer leaflet (*J_s,_*_out_). In general, increasing *J_s,_*_out_ or *R*_m,iv_ shifts the thermodynamic transition toward higher *p*_0_ values. The coexistence point *p*_0_ is highly sensitive to *J_s,_*_out_ and *R*_m,iv_, while the effect of *J_s,_*_iv_ remains negligible across the sampled parameter space (Fig. 7).

**Figure 7:**
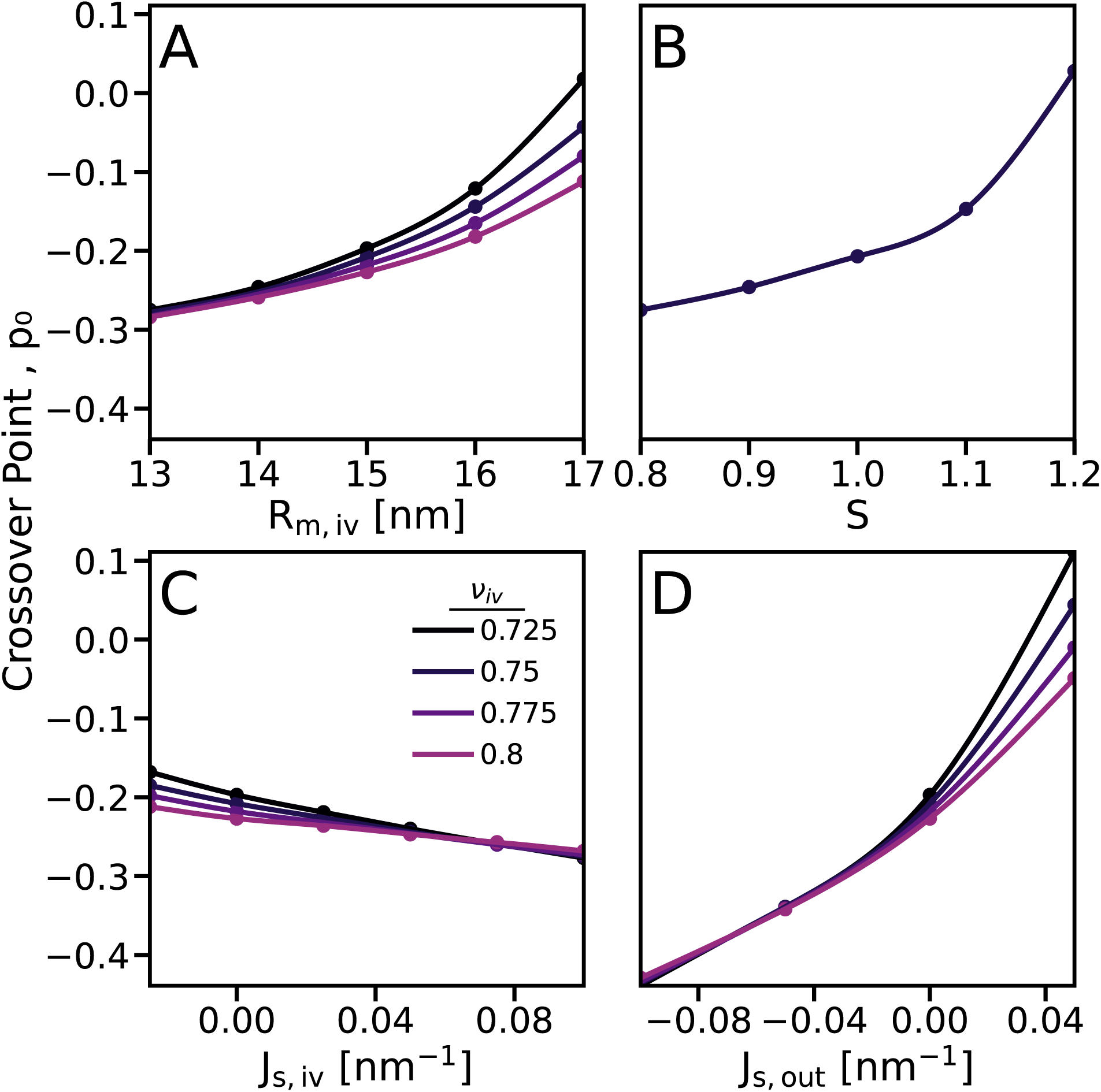
Sensitivity of the coexistence point *p*_0_ to variations in physical parameters across different *ν*_iv_. (A) Impact of the invaginating vesicle radius (*R*_m,iv_); (B) Effect of global system scaling; (C) Variation with the spontaneous curvature of the inner leaflet of the invaginating vesicle (*J_s,_*_iv_). (D) Influence of the spontaneous curvature of the shared outer leaflet (*J_s,_*_out_); Solid lines represent cubic spline interpolations.

While high *J_s,_*_out_ and *R*_m,iv_ values eliminate the eye-to-kettle barrier (Fig. 8, A-B), system scaling produces a non-monotonic response. The energy peaks at 167 k_B_T near *S* ≈ 1.6. Beyond this maximum, the barrier enters a regime of gradual decay (Fig. 8, C). Numerical convergence degrades as system size increases, requiring an *A/S* +*C* fit over the last 3 data points to characterize the asymptotic limit. This extrapolation suggests that the barrier remains finite, saturating at approximately 96 *k*_B_*T* .

**Figure 8:**
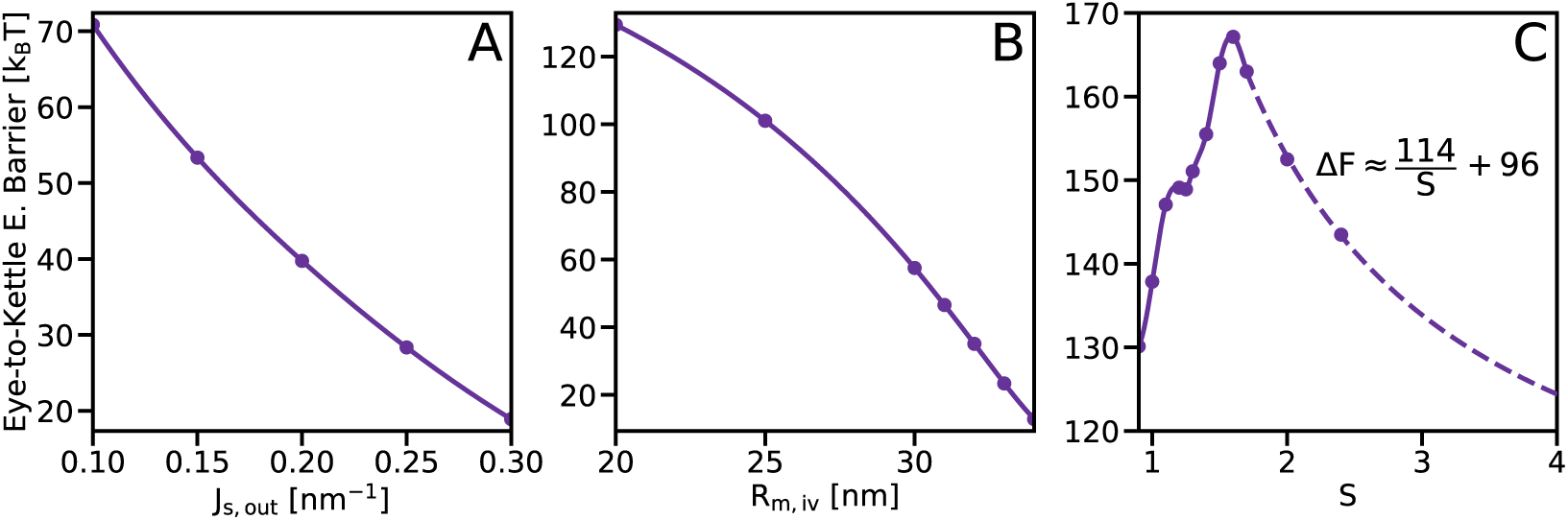
Evolution of the eye-to-kettle energy barrier at *ν*_iv_ = 0.75 and *p*_0_ = 0 across a physical parameter space. From left to right: variations in the spontaneous curvature of the shared outer leaflet (*J_s,_*_out_), the IV radius (*R*_m,iv_), and the global system scale (*S*). Solid lines denote cubic spline interpolations; the dashed line in the scaling plot indicates reciprocal extrapolation in the high-scale regime where sampling density is reduced.

## 4 Discussion

### 4.1 Snap-Through Dynamics and Energetics

When the invaginating vesicle (IV) volume is low (*ν*_iv_ ≤ 0.825), the transition between the eye and kettle morphologies manifests as a discontinuous snap-through event (Fig. 3). This transition represents an energetic trade-off between IV deformation and HV leaflet stretching. In the eye state, the IV maintains a lens-like geometry that minimizes the IV deformations but requires a high internalized volume ratio (Fig. 5). This volume displacement forces the HV leaflets to stretch significantly to accommodate the protrusion. Conversely, the kettle morphology deforms the IV into a high-curvature, elongated state with a constricted neck. This increases the IV bending penalty but reduces the internalized volume and alleviates the stretching cost on the HV. Ultimately, the system minimizes the total free energy by partitioning the elastic strain between the two vesicles according to available leaflet area and volume constraints.

Figure 6 shows that the eye-to-kettle transition at *ν*_iv_ = 0.725 is governed by a ≈ 70 *k*_B_*T* kinetic barrier at the thermodynamic crossover. The system remains kinetically locked in the eye morphology until *p*_0_ decreases sufficiently to cross the barrier, triggering an abrupt jump to the kettle state. The transition is highly hysteretic; the barrier to revert to the eye geometry remains above 30 *k*_B_*T* (Fig. 6, A) at *ν*_iv_ = 0.725. This energy floor, comparable to the known cost of a membrane fusion stalk ^6^, ensures that the IV maintains its kettle shape even if *p*_0_ increases during externalization. This abrupt transition may function as a binary mechanical switch, such that the release of stretching stress during the eye-to-kettle shift could potentially serve a physical signal to trigger protein recruitment or vesicle maturation.

### 4.2 Physical Determinants of Morphological Stability

The shared outer leaflet constitutes the majority of the membrane area. Its elastic contributions, specifically the bending penalty from positive spontaneous curvature, govern the internalization energy. Positive *J_s,_*_out_ penalizes the inward curvature required for the eye morphology’s protrusion^3,20^, which shifts the equilibrium toward the kettle state (Fig. 7, A). Similarly, increasing *R*_m,iv_ stabilizes the kettle (Fig. 7, B). Larger invaginating vesicles utilize the kettle geometry to markedly minimize the internalized volume (Fig. 5). This structural efficiency reduces the total elastic strain on the host vesicle and favors the kettle phase at higher *p*_0_.

### 4.3 Morphology in Hemifusomes

Hemifusomes are specialized vesicular organelle complexes observed in endocytic pathways, hypothesized to facilitate efficient cargo internalization^11^. Both “eye” and “kettle” morphologies have been observed in experimental cryo-electron tomography of hemifusomes. The “eye” geometry appears at higher *p*_0_ and the “kettle” at lower *p*_0_, which is consistent with our model’s predictions. Notably, the experimental micrographs suggest that internalized hemifusomes may operate above the critical point (large *ν*_iv_). For instance, the structures in Ref. ^11^ (Fig. 4b left and 4c right) closely resemble the continuous evolution that we find in our model results at *p*_0_ ≈ −0.6 and *ν*_iv_ = 0.875 (fig. 4). In this regime, the structural evolution from eye-like to kettle-like remains well-defined and occurs around the same *p*_0_ coordinate as the discontinuous transitions observed at lower *ν*_iv_ (fig. 3). The relevant physical parameters thus continue to shift this crossover location, dictating morphological preference even in the absence of a discrete jump. Cryo-ET imaging in Ref. ^11^ identified a proteolipid nanodroplet, a high-density protein-lipid condensate, localized at the hemifusion junction. This associated nanodroplet may promote spontaneous curvature in the shared outer leaflet to drive morphological selection. Beyond geometric selection, the nanodroplet likely drives the system along the *p*_0_ coordinate by acting as a localized lipid reservoir. The droplet manipulates local area asymmetry by sequestering lipids from the shared outer leaflet or releasing them into the inner leaflet. This process steers the complex toward the smoothened crossover point.

### 4.4 Scaling and Competitive Pathways

The absence of kettle morphologies in synthetic GUV experiments creates a thermodynamic-experimental paradox. While our model predicts the kettle as the dominant equilibrium state at the micron scale (Fig. 7), to our knowledge, it has not been reported in GUV studies^11,21,22^. This discrepancy likely stems from two non-exclusive factors. First, our scaling analysis suggests that the eye-to-kettle barrier does not vanish with increasing size. Instead, it saturates at an extrapolated limit of ≈ 96 *k*_B_*T* (Fig. 8, C). This provides strong evidence that GUV-scale systems are effectively kinetically trapped. The transition to the thermodynamically favored kettle morphology is thus practically inaccessible. Thermal fluctuations cannot overcome such a substantial barrier on experimental timescales. Second, our model may lack necessary dampening factors like undulatory entropic contributions or the complexity of biological membranes. These factors could alleviate the internalization stresses that result in kettle dominance in our continuum model and their exclusion in experiments.

The internalization process requires extensive leaflet area redistribution. Specifically, achieving high Φ_in_ values requires an area shift comparable in scale to the total number of lipids in the IV’s outer leaflet. This requirement far exceeds values reported for standard budding processes, where the necessary lipid asymmetry is relatively low ^3^. Such a costly process likely cannot be driven solely by localized fluctuations or small lipid sinks at the GUV scale. In those systems, the required lipid flux would exceed the capacity of a localized proteolipid complex. Consequently, internalization competes with full fusion; the energetic cost to reach the snap-through point likely exceeds the barrier for pore formation. The kettle is therefore a specialized intermediate that requires the nanometric environment of the hemifusome to bypass these kinetic traps and avoid premature fusion.

## 5 Conclusion

We characterized a morphological phase transition in LUVs that governs the internalization pathway of hemifused vesicles. This transition, which shifts the system from a lens-like “eye” to an elongated “kettle” geometry, is regulated by the reduced volume of the invaginating vesicle (*ν*_iv_). While deflated invaginating vesicles undergo a discontinuous snap-through process, those with higher relative volumes exhibit a continuous structural evolution. The kettle geometry minimizes the internalization ratio (Φ_in_) to alleviate the elastic stretching penalty on the host vesicle. The global stability of these profiles is strictly dictated by the physical parameters of the complex; specifically, increasing the spontaneous curvature of the shared outer leaflet (*J_s,_*_out_) or the invaginating vesicle radius (*R*_m,iv_) shifts the equilibrium toward the kettle morphology. Furthermore, the transition is inherently hysteretic. Once achieved, the high-curvature neck of the kettle geometry remains metastable even during externalization, preventing spontaneous reversion to the eye state and ensuring unidirectional morphological persistence throughout the internalization process.

The biophysical framework presented here provides a direct roadmap for identifying these membrane remodeling processes in vivo and replicating them in synthetic systems. Biologically, the sharp, hysteretic nature of the snap-through transition suggests that the cell can exploit the shifting global membrane elastic stresses as a mechanical checkpoint. The proteolipid nanodroplet within hemifusomes could thus function as a low-cost thermodynamic catalyst: by subtly tuning local outer-leaflet spontaneous curvature or acting as a localized lipid sink, it can drive the system across the continuous or discontinuous threshold to trigger directional cargo internalization without sustained ATP-driven remodeling. Validating these energetic predictions empirically requires capturing the nanoscale invaginating vesicle volume (*ν*_iv_). To this end, 3D cryo-electron tomography (cryo-ET) of heterogeneous vesicle populations offers the most viable path, as full volumetric reconstruction is necessary to precisely quantify vesicle volumes and surface areas. Post-facto 3D segmentation will allow concurrent calculation of both invaginating vesicle’s relative water content and internalization ratio from individual tomograms. Plotting these experimental coordinates directly onto our calculated phase diagrams will reveal whether real biological complexes cross a sharp snap-through threshold or trace a smooth morphological continuum. Ultimately, this mapping offers a powerful secondary benefit: by matching the observed intermediate shapes and their dynamic transition rates to our model’s observed structures and calculated energy barriers, researchers can work backward to implicitly extract the elastic properties of the hemifused vesicles and possibly each individual leaflet.

## Author contributions

**Itay Schachter:** Software, Investigation, Formal analysis, Writing – original draft. **Daniel Harries:** Supervision, Writing – review & editing. **Pavel Jungwirth:** Supervision, Writing – review & editing.

## Data availability

The continuum elastic model implementation and the numerical finite-element solver used in this study are open-source. The underlying code, along with the scripts and parameters required to reproduce the simulations and energy landscapes presented in this article, are freely available on GitHub at https://github.com/infinityScha/ElasticTool/tree/invaginating-vesicles. Additional processed data are available from the corresponding author upon reasonable request.

## Acknowledgements

P.J. acknowledges support from the European Research Council (ERC Advanced Grant No. 101095957). I.S. acknowledges support from Charles University and from the International Max Planck Research School for Quantum Dynamics and Control hosted by the Max Planck Institute for the Physics of Complex Systems, Dresden, Germany. Financial support to D.H. from the Israel Science Foundation (ISF Grant No. 1207/21) is gratefully acknowledged. The authors acknowledge the HPCg computational facility at the Institute of Organic Chemistry and Biochemistry (IOCB) Prague for providing computational resources. During manuscript preparation, the authors used Google Gemini for formatting, language editing and generating graphical elements within the TOC figure; the authors reviewed all outputs and take full responsibility for the final content.

