## Supplementary material for "Vesicle Internalization Proceeds via a Morphological Phase Transition": Supplimentary Information and Figures

### Supplementary Information: Vesicle Internalization Proceeds via a Morphological Phase Transition

June 15, 2026

#### S1 Numerical Discretization and Pathway Sampling

Our methodology employs a finite-element solver that utilizes a grid-based discretization of the $(r, z)$ midplane and leaflet surfaces to evaluate equilibrium structures and transition pathways, as detailed previously [1]. The core elastic energy functional follows the procedure previously described. Here, the energy functional and methodological details have been modified because the hemifused geometry requires a redefined reaction coordinate for path sampling and specialized grid refinement at the junction to accurately resolve high bending energy gradients. In the following sections we outline the modifications we have made to the methodology.

##### S1.1 Minimal Free Energy Path (MFEP) Sampling

In our original methodology, MFEP sampling followed the Partial Nudged Elastic Band (PNEB) method [1, 2], which utilized two representative grid points on the outer leaflet as anchors to determine an appropriate reaction coordinate $\xi$. Here, this was modified to accommodate the hemifused topology by redefining the reaction coordinate anchors as two points flanking the junction on the IV inner leaflet. These anchors are fixed at half the total arc-length from the junction, as measured in the kettle morphology, for each respective bilayer patch. To prevent arbitrary rigid-body translations and to establish a consistent spatial reference frame across all images, the midpoint of the central hemifusion junction profile is constrained to the coordinate origin by pinning its position strictly to $z = 0$.

The minimal free energy path (MFEP) was discretized using $M = 7$ images. The transition pathways are smooth and exhibit a single localized barrier (Fig. S2). Consequently, $M = 7$ provides an overdetermined string with 5 intermediate degrees of freedom. Prior work confirmed $M = 5$ is sufficient to resolve similar energy profiles without loss of fidelity [1]. Using $M = 7$ provides sufficient resolution such that the piece-wise cubic spline interpolation and calculated barrier heights are insensitive to further path refinement. This choice simultaneously optimizes computational throughput.

##### S1.2 Equilibrium Minimization and Sampling Strategy

To map the energy landscape and seed the MFEP calculations, initial configurations were manually constructed as two hemifused spherical vesicles. These were minimized at $\nu_{iv} = 0.75$, $\nu_{hv} = 1$, and the internalization coordinate extremes ($p_0 = -0.8$ and $p_0 = 0.4$) to seed the kettle and eye morphologies, respectively (see Fig. 2). Subsequent minimized configurations across the parameter space were calculated using continuation methods, where a previously converged solution served as the initial guess for adjacent parameter values. For system scaling ($S$), the initial structure was scaled proportionally to maintain a proximity to the global minimum, thus improving the stability of the solver during the initial minimization steps.

##### S1.3 Hemifusion Junction Refinement and Resolution Scaling

As introduced above, the hemifused geometry requires specialized grid refinement at the junction to accurately resolve high bending energy gradients. This refinement ensures that morphological transitions are physically attainable rather than resulting from discretization artifacts. We implemented a spatially-dependent resolution scheme to stabilize the solver at the hemifusion site. For the three-bilayer junction point, we apply the following discretization rules:

- **High-Resolution Zone:** The 3 grid elements closest to the junction are fixed at 0.5 nm.
- **Transition Zone:** Over the next 5 elements, grid spacing increases linearly from 0.5 nm to the global baseline resolution ($ds_m$).
- **Baseline Zone:** Standard resolution ($ds_m$) is maintained for the remainder of the vesicle body.

This moderating graded transition in element size imposes numerical stability, because abrupt resolution discontinuities destabilize the solver, and induce cubic spline interpolation kinks that can generate spurious energy minima and numerically trap the optimizer. Smoothing the discretization gradient ensures a continuous curvature representation across the junction.

The global baseline resolution of the bilayer midplane, $ds_m$, further from the junction was adapted for specific simulation tiers:

1. **Equilibrium Structure Mapping:** we used $ds_m = 0.5\,\text{nm}$ for Figures 2–5 and 7 to ensure precise crossover coordinates.
2. **Path Sampling (MFEP):** $ds_m = 1.0\,\text{nm}$ was used to maintain computational efficiency during band relaxation. We verified that this reduction shifts the total energy by less than $0.4\,k_\text{B}T$ for the studied morphologies at $\nu_{iv} = 0.75$ and $p_0 = 0$, having a negligible effect on barrier height calculations.
3. **Large-Scale Systems ($S \geq 1.2$):** For simulations involving significantly larger vesicles (e.g., Fig. 8C), $ds_m$ was increased to 1.5 nm for $S \in [1.2, 1.4)$, 2 nm for $S \in [1.4, 2)$, and 3 nm for $S \geq 2$. This adjustment was necessary to ensure numerical convergence within reasonable timescales as the total membrane area increased and, accordingly, also the total number of nodes. This coarser mesh does not significantly impair result quality because bending deformations and the rates of change in other deformation modes further from the hemifusion junction scale as $1/S$.

### S2 Supporting Figures

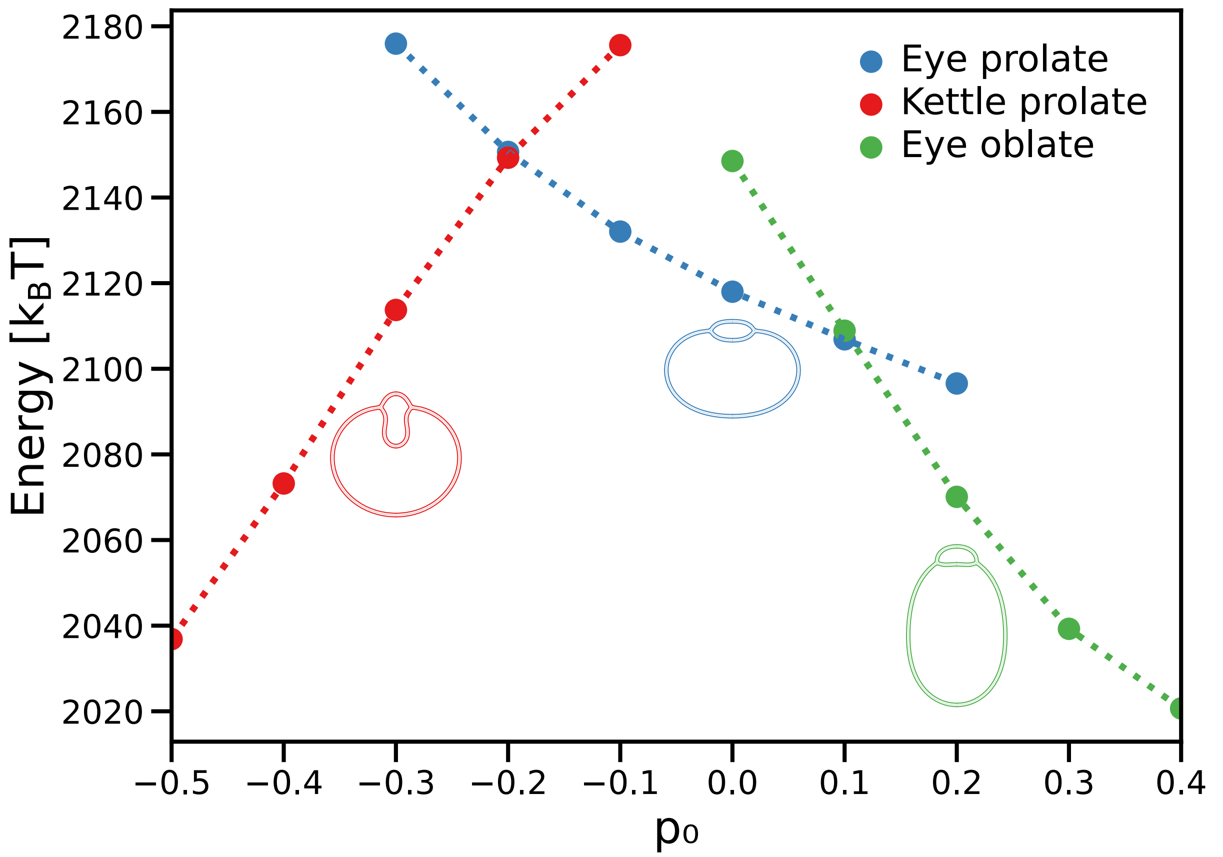


Figure S1: Energy landscape of the prolate "eye" (blue), oblate "eye" (green), and "kettle" (red) morphologies as a function of the internalization coordinate $p_0$ at $\nu_{iv} = 0.725$. Scatter points indicate discrete $p_0$ coordinates of the energy minimizations; dotted lines are linear interpolations.

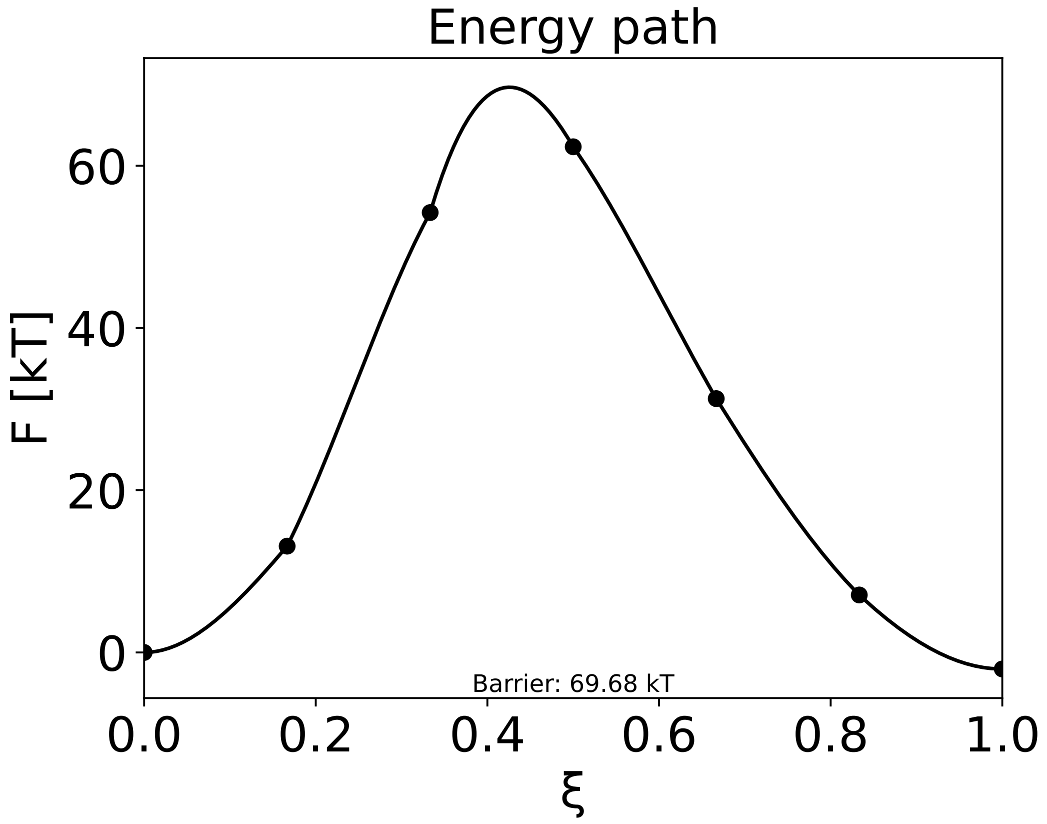


Figure S2: Example of a minimal free energy path at the internalization coordinate $p_0 = -0.2$ and $\nu_{iv} = 0.725$ over the found reaction coordinate $\xi$. Scatter points indicate discrete images energy values. Solid line is a piece-wise cubic spline derived from the energies and the parallel component to the path of the forces between the images.
